# Tissue-Specific Enrichment Analysis (TSEA) to decode tissue specificity

**DOI:** 10.1101/456293

**Authors:** Guangsheng Pei, Yulin Dai, Zhongming Zhao, Peilin Jia

## Abstract

**Motivation:** Diseases and traits are under dynamic tissue-specific regulation. However, heterogeneous tissues are often collected in biomedical studies, which reduce the power in the identification of disease-associated variants and gene expression profiles.

**Results:** We present *TSEA*, an R package to conduct Tissue-Specific Enrichment Analysis (TSEA) with two built-in reference panels. Statistical methods are developed and implemented for detecting tissue-specific genes and for enrichment test of different forms of query data. Our applications using multi-trait genome-wide association data and cancer expression data showed that TSEA could effectively identify the most relevant tissues for each query trait or sample, providing insights for future studies.

**Availability:** https://github.com/bsml320/TSEA

## 1 Introduction

Genome-wide association studies (GWAS) and next-generation sequencing technologies have identified hundreds of thousands of disease-associated variants and genes. Interpretation of these variants, however, remains an open challenge. Tissue-specific regulation, which is affected by many genetic variants, is a critical factor leading to diseases or traits. So far, many diseases or traits do not have causal tissues or cell types, or in particular with its regulation. Recent success of the Genotype-Tissue Expression (GTEx) (GTEx Consortium, 2013) enables us to systematically investigate tissue-specific gene expression and regulation. Leveraging these non-disease, reference data could open new avenues to infer causal tissues for diseases and to unveil underlying biological mechanisms.

Tissue transcriptome data is often heterogeneous. It includes both genes that are ubiquitously expressed (e.g., housekeeping genes) and genes that are specifically expressed in a range of tissues. Several methods have been developed to identify tissue-specific genes (TSGs) from expression profiles. For example, SpeCond fits a normal mixture model to each gene using microarray data for 32 human tissues (Cavalli, et al., 2011). Zhao, et al. (2015)used the Tukey test to identify TSGs from RNA-sequencing (RNA-seq) data. Here, we present a convenient R package, Tissue-Specific Enrichment Analysis (TSEA), to identify the most relevant tissues for candidate genes or for gene expression profiles. TSEA builds on two pre-processed reference panels. We developed a statistical method to identifying TSGs while controlling potential confounding factors. We implemented different statistic tests for different forms of query data. We validated the reference panels by comparing each other and demonstrated TSEA using multi-trait GWAS data and cancer RNA-seq data. Finally, we provided the methods and the reference panels in the R package, *TSEA*.

## 2 Methods

### 2.1 Data collection

We prepared two reference panels: GTEx and the Encyclopedia of DNA Elements project (ENCODE). The GTEx RNA-seq data included 14,725 protein-coding, non-housekeeping genes in 47 tissues (Eisenberg and Levanon, 2013) (details in Text S1 and Table S1). The ENCODE panel included 14,031 protein-coding, non-housekeeping genes for 44 tissues (accessed August 2018, Table S2). We downloaded and processed GWAS summary statistics for 26 traits (Table S3). We defined trait-associated genes (TAGs) by using the gene-based p-values (Text S1). In addition, we downloaded RNA-seq data for 635 normal samples matched to 14 cancers from The Cancer Genome Atlas (TCGA, Table S4).

### 2.2 Measurement of tissue specificity

For GTEx data, we implemented a previous method (Finucane, et al., 2018) by fitting an ordinary regression model for each gene and computed *t*-statistics to measure the tissue specificity. Notably, several tissues in GTEx dataset were biologically related, such as some brain sub-regions. Treating these instinctively related tissues as independent and including all of them in one regression model would underestimate the tissue-specificity. Finucane et al. (2018) defined a new variable called “tissue group” (Table S1) and used it as the explanatory variable. We followed their work and fitted the regression model for each tissue as: *Y* ~ *X* + age + sex, where *Y* was the log2-transformed gene expression, the nominal variable *X* was the tissue group status, and age and sex were sample covariates. We fitted the above model for each tissue, instead of fitting one model including all tissues. Specifically, for a tissue in examination, we defined *X* = {*x_i_*}, *i* = 1,…, *N*, where *N* is the total number of samples, *x_i_* = 1 if the sample belonged to the tissue in examination and *x_i_* = 0 if the sample belonged to any tissues not in the same group. Samples from other tissues of the same group as the examined tissue would not be included. Accordingly, *N* varies by the tissues in examination. After fitting the model, we selected the *t*-statistic for the explanatory variable *X* for the gene.

Considering that sample size per tissue is small in ENCODE dataset, we employ *z*-score to measure tissue specificity. For each gene, a *z*-score is calculated as *z_i_* = (*e_i_* – mean(*E*))/*sd*(*E*), where *ei* is the average expression of the gene in the *i*^th^ tissue, *E* represents the collection of its average expression in all tissues, and *sd* indicated the standard deviation of *E*.

A higher *t*-statistic or *z*-score indicates that the gene is more specifically expressed in the corresponding tissue.

### 2.3 Tissue-specific enrichment analysis

For each tissue, TSGs are defined by high *t*-statistics or *z*-scores. We allow the user to define the cutoff values, e.g., the top 5% genes as TSGs. Depending on the query data, two tests are implemented.

Test 1: if the query is a list of genes, we implement Fisher’s Exact Test to identify TSGs enriched in the tissue(s).

Test 2: if the query is an expression matrix, we use *t*-test to identify the most relevant tissue(s). Two methods are used to normalize the query expression data so that the query data will be scaled appropriately with the reference data. (1) The *z*-score strategy that normalizes the query data using the tissue parameters as below: *e_n_* = (*e_q_*-*u_s_*)/*sd_s_*, where *e_q_* and *e_n_* are the query and normalized expression, and *u_s_* and *sd_s_* are the mean and *sd* of a reference tissue *s*. (2) The abundance correction approach (Skene, et al., 2018) that normalizes the query data by *e_n_* = *log_2_*(*e_q_*+1)/(*log_2_* (*u_s_*+1)+1). Finally, two-sample *t*-test is used to examine the difference between *e*n of TSGs and *e*_n_ of non-TSGs in each reference tissue (Fig. S1).

## 3 Applications

We provided two reference panels in *TSEA*: the GTEx panel (47 tissues, *t*-statistic) and the ENCODE panel (44 tissues, *z*-score). Validation across the two panels showed high concordance (91.5 – 93.2%, Text S1). TSEA can be applied in many scenarios, such as inferring the causal tissues for diseases based on disease-associated genes, assessing bulk RNA-seq data (e.g., finding the most related tissues, outliers, sample contamination/purity), and cross-validation of other genomics data (e.g., tissue-specificity of microRNAs or transcription factors by their targets), among others. Below we demonstrate the utility of TSEA with two applications.

### Application 1: multi-trait GWAS data

We tested TAGs (gene-based p < 1×10^−6^) identified from GWAS for 26 traits using TSEA Test 1 for candidate genes. In most traits, TAGs were found enriched in the trait-related tissues whereas instances of variation implied novel insights into the disease origin(s). For example, TAGs of anthropometric traits were mainly enriched in artery tissues, metabolic traits in liver, and neurologic/social traits in brain. However, autism spectrum disorder, body mass index, and fasting glucose failed to be linked with any tissues, likely due to weak GWAS signals or the causal tissues not included in our panels.

### Application 2: TCGA normal samples

The RNA-seq data for TCGA normal samples were organized as an expression matrix and were analyzed using TSEA Test 2. For demonstration purpose, we pre-defined the biologically matched tissue of each cancer type (Text S1, Table S4). We normalized the original RNA-seq data using the abundance correction strategy based on the GTEx panel. As a result, in nine cancer types (denoted by triangle in Fig. 1B), the matched tissues were most enriched in the samples. In four cancer types (circles in Fig. 1B), the biological tissues were ranked within top 3 and the related tissues could be implied. For example, stomach was the most enriched tissue for esophagus cancer, providing insights into cancer origination. In breast cancer, only 50% samples were most enriched in breast while others in minor salivary gland and uterus. This result is likely due to sample purity.

**Fig. 1.**
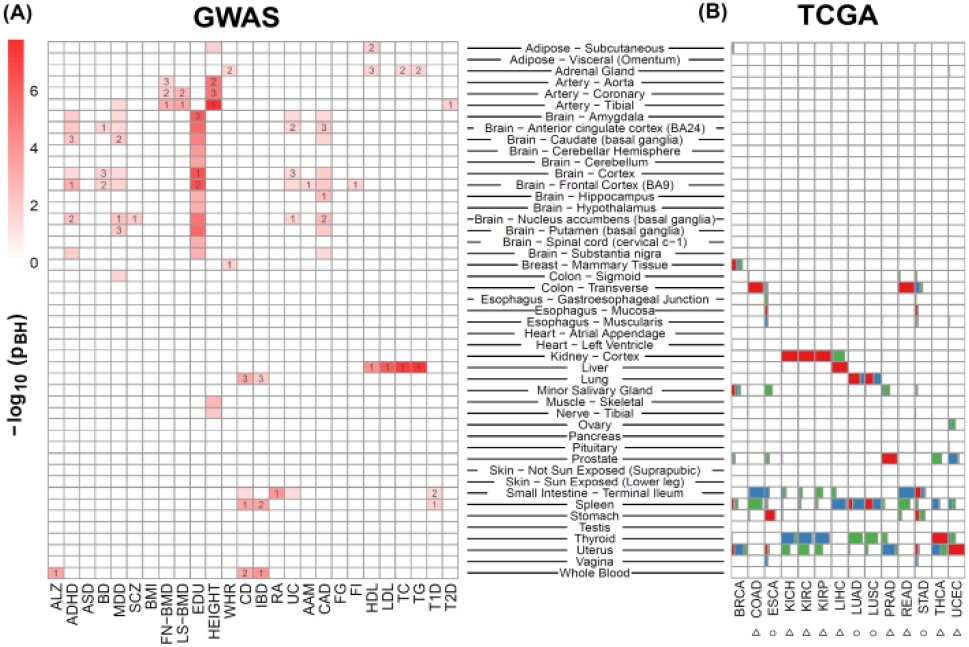
Tissue-specific enrichment analysis of GWAS and TCGA data. Full names of traits and cancer types are available in Tables S3 and S4. The p-value (adjusted by the Benjamini and Hochberg method) was from Fisher’s Exact Test (A) and *t*-test (B), respectively. In (B), boxes in red (top 1), blue (top 2), and green (top 3) indicate the proportion of samples enriched in a tissue.

## 4 Conclusion

We presented an R package, *TSEA*, for tissue-specific enrichment analysis. *TSEA* runs fast – it took 2 minutes for a RNA-seq matrix with 635 samples on an i7-7700HQ desktop. *TSEA* is useful to study tissue features and underlying mechanisms for diseases or traits.

## References

Cavalli, F.M.etal. SpeCond: a method to detect condition-specific gene expression. Genome Biol 2011;12(10):R101.

Eisenberg, E. and Levanon, E.Y. Human housekeeping genes, revisited. Trends in genetics: TIG 2013;29(10):569-574.

Finucane, H.K.etal. Heritability enrichment of specifically expressed genes identifies disease-relevant tissues and cell types. Nature genetics 2018;50(4):621-629.

GTEx Consortium. The Genotype-Tissue Expression (GTEx) project. Nature genetics 2013;45(6):580-585.

Skene, N.G.etal. Genetic identification of brain cell types underlying schizophrenia. Nature genetics 2018;50(6):825-833.

Zhao, G.etal. An effective analytic method for detecting tissue-specific genes in RNA-seq experiments. Pharmacogenomics 2015;16(16):1769-1779.

